# ProtFun: A Protein Function Prediction Model Using Graph Attention Networks with a Protein Large Language Model

**DOI:** 10.1101/2025.05.13.653854

**Authors:** Muhammed Talo, Serdar Bozdag

## Abstract

Understanding protein functions facilitates the identification of the underlying causes of many diseases and guides the research for discovering new therapeutic targets and medications. With the advancement of high throughput technologies, obtaining novel protein sequences has been a routine process. However, determining protein functions experimentally is cost- and labor-prohibitive. Therefore, it is crucial to develop computational methods for automatic protein function prediction. In this study, we propose a multi-modal deep learning architecture called ProtFun to predict protein functions. ProtFun integrates protein large language model (LLM) embeddings as node features in a protein family network. Employing graph attention networks (GAT) on this protein family network, ProtFun learns protein embeddings, which are integrated with protein signature representations from InterPro to train a protein function prediction model. We evaluated our architecture using three benchmark datasets. Our results showed that our proposed approach outperformed current state-of-the-art methods for most cases. An ablation study also highlighted the importance of different components of ProtFun. The data and source code of ProtFun is available at https://github.com/bozdaglab/ProtFun under Creative Commons Attribution Non Commercial 4.0 International Public License.

## 1 Introduction

Proteins, the fundamental building blocks of all living systems, are involved in a wide range of essential functions in cells and play a crucial role in cell structure, facilitate chemical reactions, and transmit signals from the external environment. Understanding the functions of proteins is essential for understanding the biological processes that operate inside of an organism (1).

The role that a protein plays in an organism is determined by its function. Predicting protein function is a multi-label classification task. Protein functions are generally described with gene ontology (GO) terminology, which has above 41,000 terms as of February 2025. GO terminology is organized into three categories, namely molecular function ontology (MFO), biological process ontology (BPO), and cellular component ontology (CCO). All these subontologies have a directed acyclic graph (DAG) structure. While the more specialized protein activities are found at leaf nodes, the more general protein functions are outlined in the parent nodes. A protein may have numerous distinct GO terms. The time-consuming and expensive experimental techniques are used to discover protein functions. The GO database has experimental annotations for only one percent of proteins (2). Therefore, it has become indispensable for researchers to create computational methods for the automatic prediction of protein function.

A wide range of data types have been used to predict protein functions, including protein sequence (3), structure (4), protein-protein interaction (PPI) (5), phylogenetic information (6) and literature information (5). It is crucial to be aware that using literature knowledge can be beneficial, but it must be managed cautiously to prevent information leakage into test set, which could inflate performance metrics spuriously. In addition, rather than relying on a particular data type, substantial advancements have been achieved by integrating diverse data types using integrative approaches to identify protein functions (5–7).

Sequence alignment and domain based analysis are fundamental approaches in protein function prediction. Tools such as DIAMOND (8), BLAST (9), and InterProScan (10, 11) are commonly used for predicting protein functions. For example, Kulmanov and Hoehndorf employed BLAST and DIAMOND sequence alignment algorithms to infer the functions of proteins (3) based on the assumption that proteins with similar sequences have similar functions. However, it is also possible for proteins with different sequences to have the same functions (12). InterProScan facilitates the identification of protein families, domains, and sites by integrating information from 14 distinct databases. It generates signatures by comparing protein sequences with diverse patterns and categorizes proteins into families and predict domain presence using similarity search algorithms. The signatures encapsulate the predefined pattern and motif information. InterProScan has been utilized as an effective tool for depicting diverse protein features in several studies (13, 14).

Around 80% of proteins commonly carry out their functions through interactions with other proteins (15). To utilize this information, graph neural networks (GNN)-based methods have been employed on PPI networks to enhance protein function prediction (14). However, the limited availability of PPI in biological systems, as compared to protein sequences, emphasizes the limitations of this method. For instance, in DeepGraphGO, You et al. (14) was able to use only 65% of the training data by conducting training on 17 specific organisms.

The application of language modeling approaches using protein sequences has enhanced the precision of protein function detection. Recurrent neural networks (RNNs), 1D convolutional neural networks (CNNs), and large language models (LLMs) pre-trained on protein sequences are among the most frequently employed models. For example, Kulmanov and Hoehndorf used a 1D CNN architecture (DeepGOCNN) in the DeepGOPlus study to predict protein functions (3). In their subsequent studies, the authors employed the ESM2 protein LLM (16) along with GO axioms to predict protein functions (17). In another study, Cao and Shen developed a transformer-based protein function annotation through joint sequence–label embedding (TALE) model utilizing selfattention-based transformers to capture global patterns in protein sequences and embed hierarchical GO annotations into a joint latent space (18).

Many machine learning and bioinformatics teams around the world have been inspired to develop computational techniques to participate in the Critical Assessment of Function Annotations (CAFA) protein function prediction competition (19). In the last decade, CAFA contests have resulted in significant advancements in the prediction of protein functions by fostering the integration of traditional methods with state-of-the-art (SOTA) deep learning methods. For instance, in the CAFA3 competition, GOLabeler (20) achieved the best performance for protein function prediction by employing a learning-to-rank (LTR) framework that integrates five component classifiers trained on multiple feature types, including GO term frequency, sequence alignment, amino acid trigrams, domains and motifs, and biophysical parameters. The most recent version of the NetGO model has demonstrated its effectiveness in both CAFA4 and CAFA5 contests. In CAFA4, NetGO model utilized a protein LLM with literature information, while in CAFA5, it further integrated additional data types, such as protein textual descriptions, protein 3D structure, and scientific literature, to enhance protein function prediction (7).

There are several limitations of the existing protein function prediction methods. Sequence-similarity based methods are unable to capture the complex patterns in the sequence. GNN-based methods on PPI networks provide limited generalizability as PPI information is available for only a limited number of well-studied organisms. Protein LLM-based methods have had high accuracy, but they only utilize sequence data.

To address these limitations, in this study, we propose a framework called ProtFun that employs graph attention network to integrate LLM-based embeddings and InterPro signatures on protein domain and family information. In order to demonstrate the effectiveness of our proposed model, we conducted model training and evaluation on three benchmark datasets. Our proposed architecture exhibits state-of-the-art (SOTA) performance in nearly all cases in BPO, MFO, and CCO term predictions in terms of the CAFA metrics, *F*_max_ and *S*_min_, as well as the area under the precision-recall curve (AUPRC). The source code, documentation, and output of the ProtFun model are available as Jupyter Notebooks to facilitate reproducibility and can be accessed at https://github.com/bozdaglab/ProtFun.

## 2 Materials and methods

### A. Datasets

In this study, we evaluated the performance of the proposed model and compared it with the SOTA methods using three different benchmark datasets introduced in the NetGO (5), DeepGOZero (13), and DeepGraphGO (14) papers. These datasets will be referred to as the NetGO, DeepGOZero, and DeepGraphGO datasets throughout this paper. All three datasets were split into training, validation, and test sets by the dataset curators, including protein sequences and GO terms from all subontologies. To ensure a fair comparison with existing methods, we utilized the same training and test splits as those used in prior research. The distribution of the number of proteins utilized in these datasets is given in Table 1.

**Table 1.** The distribution of the number of proteins in NetGO, DeepGOZero, and DeepGraphGO datasets across subontologies.

| Dataset | Train |  |  | Validation |  |  | Test |  |  |
| --- | --- | --- | --- | --- | --- | --- | --- | --- | --- |
|  | BPO | CCO | MFO | BPO | CCO | MFO | BPO | CCO | MFO |
| NetGO | 89,828 | 81,377 | 62,646 | 1,124 | 1,359 | 1,128 | 491 | 268 | 505 |
| DeepGOZero | 47,733 | 48,318 | 34,716 | 5,552 | 4,970 | 3,851 | 5,444 | 5,969 | 4,712 |
| DeepGraphGO | 85,104 | 76,098 | 51,549 | 1,570 | 923 | 490 | 925 | 1,224 | 426 |

The NetGO dataset was created following the CAFA procedure. The train, validation, and test sets were separated based on experimentally-annotated data collected at different time intervals. The train dataset consists of proteins annotated up until December 2018. The proteins in the validation dataset were annotated between January 2019 and January 2020, while the proteins in the test dataset were annotated from February 2020 to October 2020. Protein sequences have been obtained from UniProt (21), and experimental annotations were gathered from SwissProt (22), GOA (23), and GO (2) databases. The annotations were generated using GO terms supported by experiment (EXP), direct assay (IDA), mutant phenotype (IMP), physical interaction (IPI), genetic interaction (IGI), expression pattern (IEP), traceable author statement (TAS), and curator (IC) evidence codes. Similarly, the DeepGraphGO benchmark dataset includes includes 117,170 proteins from 2,353 species, experimentally verified GO annotations, and employs CAFA-style chronological splits to enable realistic benchmarking.

The DeepGOZero dataset was generated based on the sequence similarity of proteins computed using DIAMOND. Proteins that have a similarity score of more than 50% were grouped together and 81% were used for the training, 9% for the validation, and 10% for the testing. DeepGOZero dataset includes additional evidence codes such as high throughput experiment (HTP), high throughput direct assay (HDA), high throughput mutant phenotype (HMP), throughput genetic interaction (HGI), and expression pattern (HEP), in addition to the experimental annotation evidence codes found in the NetGO dataset. To ensure statistically robust choice of labels for the model, each GO term was chosen to be associated with a minimum of 10 proteins. The number of labels used in the BPO, CCO and MFO categories of the NetGO, DeepGOZero, and DeepGraphGO datasets are given in Table 2.

**Table 2.** The number of GO term labels used in the NetGO, DeepGOZero, and DeepGraphGO datasets across subontologies.

| Dataset | BPO | CCO | MFO |
| --- | --- | --- | --- |
| NetGO | 11,192 | 1,446 | 2,329 |
| DeepGOZero | 10,284 | 1,338 | 2,065 |
| DeepGraphGO | 10,624 | 1,361 | 2,228 |

### B. The proposed ProtFun model architecture

We have developed a predictive model called ProtFun for protein function prediction. ProtFun performs late integration of two embeddings to predict protein function. The first embedding is obtained from a protein-protein similarity graph using graph attention networks (GAT), and the second embedding is obtained from a latent representation of protein signatures in the InterPro database. The flow diagram of the proposed architecture is shown in Figure 1.

**Fig. 1.**
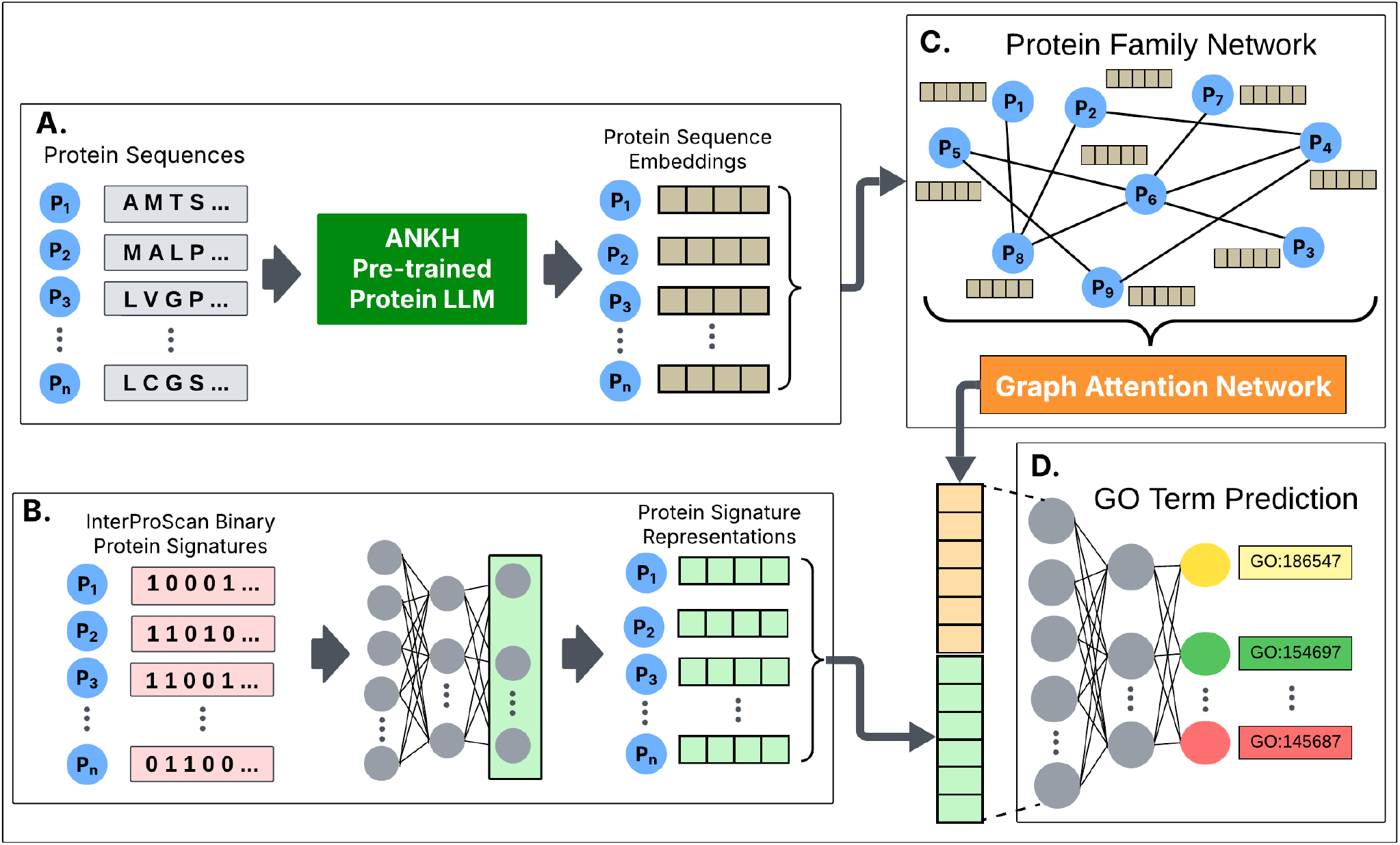
The flow chart of the ProtFun model: A) Protein embeddings were generated using ANKH LLM using protein sequences, B) Binary protein vector representations were generated based on InterPro signatures for each protein. The binary representation vector for each protein was fed into a multi-layer perceptron (MLP) model to reduce the dimensionality, C) Protein family network (PFN) was constructed using protein domain information. The protein sequence embeddings computed in (A) were used as node features of the PFN. The graph is fed into GAT to obtain the node (protein) embeddings, D) The embeddings obtained in (B) and (C) were concatenated and fed into an MLP to predict protein functions.

### C. Building protein family network from InterProScan signatures and LLM embeddings

In this study, we utilized InterProScan tool (11) and ANKH protein LLM to construct a protein family network (PFN). InterPro combines protein domains, motifs, active sites, functional regions, family data, and numerous other information to offer an understanding of the functional and structural features of proteins. Therefore, the data obtained from InterPro play a significant role for the detection of protein functions. The InterProScan tool incorporates protein signatures from a total of 14 distinct databases, namely Pfam, CDD, COILS, Gene3D, HAMAP, MOBIDB, PANTHER, PIRSF, PRINTS, PROSITE, SFLD, SMART, SUPERFAMILY, and NCBIFAM, where each protein signature represents a specific feature about the characteristics of a protein (24).

To build the PFN, we leveraged InterProScan to extract domain, motif, and family-related information. Proteins that are members of the same family, possess the same motif, or are components of the same domain were linked together by an edge to form the PFN. This network includes all potential pairs of proteins, utilizing distinctive signatures that are specifically designed to represent the relationships and interactions among proteins belonging to the same family. By structuring PFN in this manner, we categorized proteins into clusters based on their family, domain, and motif information.

We utilized the ANKH protein LLM to generate high-quality representation vectors of protein sequences, which served as node features in the PFN. ANKH utilizes a self-supervised learning approach, enabling it to learn from protein sequences without the need for explicit labels. The ANKH model was pre-trained with an encoder-decoder architecture on UniRef50 protein sequences using Google’s TPU-v4 infrastructure. The encoder maps input protein sequences into a latent embedding space, encapsulating significant sequence representations, while the decoder reconstructs the sequences by predicting the masked tokens based on the learned information. It employs a 1-gram random token masking technique, in which 15% of the sequence tokens are randomly masked and completely rebuilt. ANKH distinguishes itself by achieving high performance while minimizing computational resources because of the utilization of protein-specific experimentation instead of simply increasing the depth of the model (25). We obtained 1536-dimensional protein embeddings from the last hidden layer of the encoder, applying mean pooling across tokens to produce a fixed-size representation for each sequence.

A two-layer GNN with a GAT architecture was applied to learn enrich protein representations. The model utilizes ANKH embeddings as node features and the PFN to determine the connections among proteins. The first GAT layer was defined by 3 attention heads with a hidden dimension of 128, while the second GAT layer utilized a single head with an output dimension of 512.

### D. Latent Representation of Protein Signatures

Inter-Pro signatures were used to generate binary protein representation vectors, where a value of 1 indicates the presence of a specific InterPro signature in a protein. We utilized an MLP model to generate a latent representation of these protein signatures. The high-dimensional binary vectors passed through three fully connected layers with LeakyReLU activations to gradually reduce the dimensionality to 512, allowing effective modeling of proteins and reducing model complexity.

It is important to note that we have built the PFN, creating links among proteins having common features using InterPro signatures. Next, we created a binary representation vector for each protein, based on these signatures, to indicate the presence or absence of motifs. After that, the embeddings from GNN combined with the embeddings from the binary vector to form a new integrated representation. Aggregating PFN and binary protein representations in the same architecture is a significant design strategy, particularly when considering residual connections in deep architectures (26).

Residual connections are employed in deep architectures to mitigate information loss and enhance the learning process. The PFN offers a graphical depiction of the connections that enhance the model’s comprehension of the functional context of proteins. Binary protein representations, in contrast, quantitatively encode the specific characteristics of each protein. By combining these two data types, the proposed model is able to simultaneously access and incorporate both forms of information. By considering both protein relations and specific characteristics, the model generates more comprehensive and precise predictions for protein functions.

Finally, the graph embeddings derived from the GAT were concatenated with the latent representations of binary protein vectors and then fed into the final MLP model to predict protein function.

#### D.1 Model training and evaluation

We utilized the complete datasets of DeepGOZero, NetGO, and DeepGraphGO for our study. DeepGOZero comprises approximately 59,000 unique proteins from 1,803 species, NetGO includes around 62,000 proteins from 2,240 species, and DeepGraphGO contains 117,000 proteins across 2,353 species. This led to a graph structure that is highly dense, with an average node degree of 500 and approximately 45 million edges in the case of MFO. Therefore, we used a neighborhood sampling approach to minimize memory consumption while maintaining sufficient data for training. To effectively manage the largescale graph, a defined number of neighbors are chosen for each node at the first hop, and a smaller subset is chosen for the second hop.

The Adam optimizer and a learning rate scheduler were employed to ensure convergence and prevent overfitting. The methodology of early stopping was utilized, which involved stopping the training process if there was no improvement for a consecutive period of 5 epochs. Due to their different characteristics of subontologies, the model was trained separately for predictions related to BPO, MFO, and CCO. Training was carried out using the PyTorch library (27) and the PyTorch Geometric library (28) for graph-based operations. The model was trained and tested on a GPU server with 4 NVIDIA A100-SXM4-80GB GPUs. After the training process, the model was evaluated using an independent test set provided by the dataset creators.

### E. Metrics for evaluating model performance

To evaluate the model performance, we utilized the official CAFA-evaluator software (29). The performance was evaluated using several CAFA assessment metrics, namely *F*_max_ score, Area Under the Precision Recall Curve (AUPRC), and *S*_min_. The *F*_max_ measure is obtained using precision pr(*τ*) and recall rc(*τ*) as follows:

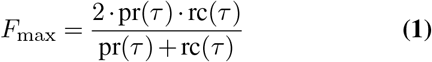

where *τ* is the cut-off value for prediction (i.e., if a protein’s score for a protein function term is ≥ *τ*, then the protein is predicted to be annotated with that term). In Equation (1), precision and recall are defined as follows:

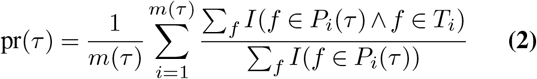

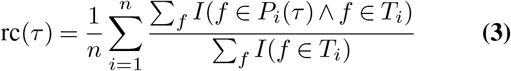

where *P*_*i*_(*τ*) is the set of predicted annotations for the cutoff *τ* for protein *i, T*_*i*_ is the set of ground truth annotations for protein *i, n* is the total number of proteins, and *m*(*τ*) is the number of predicted proteins that belong to at least one class. The identity function *I* returns 1 if the condition holds; otherwise, it returns 0. Intuitively, the highest *F* score that can be achieved for a choice of *τ* is reported as the *F*_max_ score.

*S*_min_ calculates the semantic difference between actual and expected annotations by utilizing the information content of the classes. The information content *IC*(*x*) is computed based on the annotation likelihood of class *x*. Let *P* (*x*) represent the parent classes of class *x*, then the information content of class *x* is defined as:

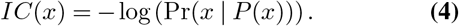

Let run(*τ*) denote the mean remaining uncertainty and mis(*τ*) denote the misinformation, calculated as:

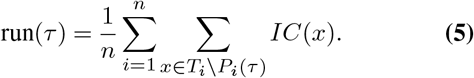

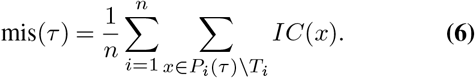

Then, *S*_min_ is defined using run(*τ*) and mis(*τ*) as follows:

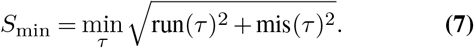

## 3 Results

We evaluated the performance of our proposed protein function prediction model, ProtFun, by comparing it with both baseline and SOTA methods. For this purpose, we adopted the evaluation scores reported in (13) and (30). Furthermore, to ensure a fair comparison, we employed the same data splits as those used in the previous studies.

### A. Evaluation on the DeepGOZero dataset

The results on the DeepGOZero dataset show that ProtFun model outperformed the SOTA models in all MFO, BPO, and CCO categories based on *F*_max_ and *S*_min_ (Table 3). For AUPRC mertic, our model outperformed all the SOTA models except for CCO where DeepGOZero (13) achieved the highest performance. The results show that our proposed model has strong ability to accurately identify hierarchical protein annotations in all three domains.

**Table 3.**
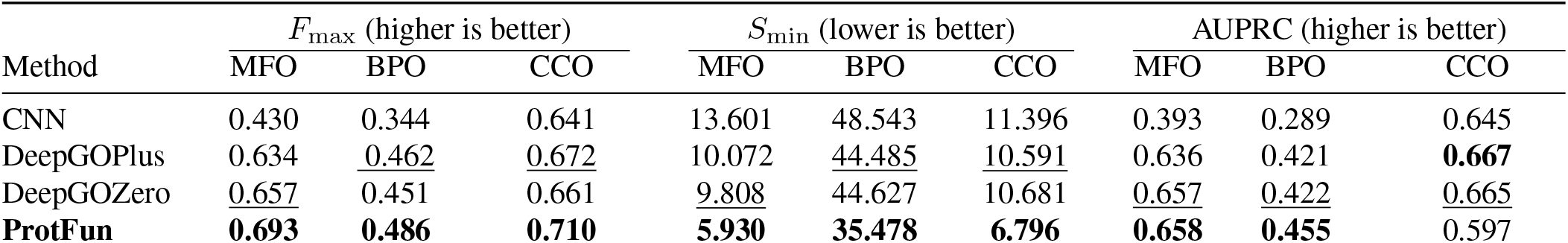
Performance of the proposed model on the **DeepGOZero** dataset. The best-performing metrics are shown in **bold** and the second-best scores are <u>underlined</u>.

| Method | $F_{\max}$ (higher is better) | | | $S_{\min}$ (lower is better) | | | AUPRC (higher is better) | | |
| --- | --- | --- | --- | --- | --- | --- | --- | --- | --- |
|  | MFO | BPO | CCO | MFO | BPO | CCO | MFO | BPO | CCO |
| CNN | 0.430 | 0.344 | 0.641 | 13.601 | 48.543 | 11.396 | 0.393 | 0.289 | 0.645 |
| DeepGOPlus | 0.634 | <u>0.462</u> | <u>0.672</u> | 10.072 | <u>44.485</u> | <u>10.591</u> | 0.636 | 0.421 | **0.667** |
| DeepGOZero | <u>0.657</u> | 0.451 | 0.661 | <u>9.808</u> | 44.627 | 10.681 | <u>0.657</u> | <u>0.422</u> | <u>0.665</u> |
| **ProtFun** | **0.693** | **0.486** | **0.710** | **5.930** | **35.478** | **6.796** | **0.658** | **0.455** | 0.597 |

In addition to evaluating the overall performance of the ProtFun model across all 1,803 species, we conducted a species-specific analysis on DeepGOZero dataset, focusing on human (taxon id: 9606), mouse (taxon id: 10090), and *Arabidopsis thaliana* (taxon id: 3702). These species were selected due to their biological significance as both mouse and *Arabidopsis thaliana* are model organisms for animals and plants, respectively. ProtFun demonstrated superior performance for *Arabidopsis thaliana*, achieving an *F*_max_ of 0.782 for MFO, 0.461 for BPO, and 0.750 for CCO indicating greater precision and reduced annotation uncertainty in plant proteins compared to the overall model performance across species (Table S1). For mouse, ProtFun achieved 0.673 (MFO), 0.449 (BPO), and 0.691 (CCO), which were slightly lower than values for across-species case. On the other hand, for human, ProtFun yielded higher *F*_max_ scores than *F*_max_ scores in across-species case for BPO and CCO.

### B. Evaluation on the NetGO dataset

The ProtFun model achieved the best *F*_max_ and *S*_min_ values for all subontologies on the NetGO dataset with the exception of the *S*_*min*_ for BPO (Table 4). In terms of AUPRC, ProtFun outperformed other models in all subontologies except for CCO.

**Table 4.** Performance of the proposed model on the **NetGO** dataset. The best-performing metrics are shown in **bold** and the second-best scores are <u>underlined</u>.

| Method | $F_{\max}$ (Higher is better) | | | $S_{\min}$ (Lower is better) | | | AUPRC (Higher is better) | | |
| --- | --- | --- | --- | --- | --- | --- | --- | --- | --- |
|  | MFO | BPO | CCO | MFO | BPO | CCO | MFO | BPO | CCO |
| DiamondScore | 0.627 | 0.407 | 0.625 | 5.503 | 25.918 | 9.351 | 0.427 | 0.272 | 0.412 |
| CNN | 0.589 | 0.337 | 0.624 | 6.417 | 27.235 | 10.617 | 0.565 | 0.271 | 0.623 |
| DeepGOPlus | 0.661 | 0.419 | 0.655 | 5.407 | <u>25.603</u> | 9.374 | 0.667 | 0.342 | 0.663 |
| DeepGOZero | 0.662 | 0.396 | 0.662 | 5.322 | 25.838 | 9.834 | 0.668 | 0.337 | 0.645 |
| NetGO2 (Webserver) | <u>0.698</u> | <u>0.431</u> | 0.662 | <u>5.187</u> | **25.076** | 9.473 | <u>0.701</u> | 0.343 | 0.627 |
| DeepGraphGO | 0.671 | 0.418 | <u>0.679</u> | 5.374 | 25.866 | <u>9.165</u> | 0.647 | <u>0.364</u> | <u>0.669</u> |
| TALE+ | 0.466 | 0.382 | 0.661 | 8.136 | 26.308 | 9.599 | 0.441 | 0.310 | **0.681** |
| **ProtFun** | **0.709** | **0.441** | **0.695** | **3.894** | 26.388 | **6.829** | **0.707** | **0.377** | 0.590 |

We have further evaluated the performance of the proposed ProtFun model on eleven representative organisms from the CAFA benchmark to examine its performance across various species. The performance of the ProtFun model for CAFA targets on the NetGO dataset is provided in Table S2. In the NetGO dataset for the CAFA targets, the ProtFun model achieved the best *F*_max_ scores of 0.799 (MFO) and 0.696 (CCO), for the human species and best *F*_max_ score of 0.532 (BPO) for fruit fly (taxon id: 7227). The precision and recall curves of the ProtFun models for human are presented in Figure S1. Our approach achieved an effective balance between precision and recall, particularly with the MFO demonstrating superior performance.

### C. Evaluation on the DeepGraphGO dataset

On the DeepGraphGO dataset, we compared ProtFun with eight baseline methods grouped into two categories: sequencebased approaches (i.e., DeepGOPlus, BLAST-KNN (20), DeepGOCNN (3), LR-InterPro (20), and PO2GO (31)) and network-based methods (i.e., Net-KNN (5), DeepGraphGO, and SEGT-GO (30)). ProtFun demonstrated improved or comparable performance across all three GO subontologies and consistently outperformed the SOTA method, SEGT-GO in *F*_max_ (Table 5).

**Table 5.** Performance of the proposed model on the **DeepGraphGO** dataset. The best-performing metrics are shown in **bold** and the second-best scores are <u>underlined</u>.

| Method | $F_{\max}$ (Higher is better) | | | AUPRC (Higher is better) | | |
| --- | --- | --- | --- | --- | --- | --- |
|  | MFO | BPO | CCO | MFO | BPO | CCO |
| BLAST-KNN | 0.590 | 0.274 | 0.650 | 0.455 | 0.113 | 0.570 |
| LR-InterPro | 0.617 | 0.278 | 0.661 | 0.530 | 0.144 | 0.672 |
| Net-KNN | 0.426 | 0.305 | 0.667 | 0.276 | 0.157 | 0.641 |
| DeepGOCNN | 0.434 | 0.248 | 0.632 | 0.306 | 0.101 | 0.573 |
| DeepGOPlus | 0.593 | 0.290 | 0.672 | 0.398 | 0.108 | 0.595 |
| PO2GO | 0.506 | 0.290 | 0.596 | 0.380 | 0.179 | 0.587 |
| DeepGraphGO | <u>0.623</u> | 0.327 | <u>0.692</u> | <u>0.543</u> | 0.194 | <u>0.695</u> |
| SEGT-GO | 0.619 | <u>0.328</u> | 0.683 | **0.555** | <u>0.217</u> | **0.703** |
| **ProtFun** | **0.678** | **0.350** | **0.719** | 0.539 | **0.258** | 0.682 |

### D. Ablation study

We performed an ablation study to evaluate the contributions of different components of ProtFun to predict MFO, BPO, and CCO terms in DeepGOZero dataset. We evaluated five distinct ablation models to examine their effect on protein function prediction as follows:

- **LLM**: This model employs only the ANKH protein LLM embeddings, removing binary protein representations and GNN features. This setting evaluates the significance of sequence context representation.
- **BPV**: This model only utilizes binary protein vectors derived from InterPro signatures, removing LLM embeddings and GNN features. It enables us to evaluate the effect of family, motif, and domain information.
- **BPV + GNN**: This ablation model uses only the binary protein vectors derived from InterProScan as input to the GNN, excluding the LLM embeddings. It evaluates the effect of using LLM as initial node features in PFN.
- **LLM + BPV**: In this model, predictions are made using the concatenation of LLM embeddings and binary protein representations skipping the GNN step. This ablation model evaluates the individual contributions of sequences and domain representations by eliminating the graph structure.
- **ProtFun (proposed method)**: This model integrates both LLM embeddings and binary protein representations, along with graph-based features obtained through GAT.

The results show that the proposed method outperformed all the ablation models in all cases except for AUPRC for CCO and MFO (Table 6). The second best model was LLM+BPV for MFO and BPO, which suggest that integration of InterPro protein structures and LLM embeddings are more useful then using either of them alone. The BPV + GNN ablation model achieves similar *F*_max_ values compared to BPV but shows better results in *S*_min_ and AUPRC across all subontologies. On the other hand, LLM + BPV outperformed BPV + GNN indicating the importance of LLM-based embeddings even in the absence of GNN. For CCO, on the other hand, LLM-based method was the second highest *F*_max_ after the proposed method suggesting that sequence-based features provide crucial information about localization in cellular structures.

**Table 6.** Performance of the ProtFun model components on the DeepGOZero dataset. The best-performing metrics are shown in **bold** and the second-best scores are <u>underlined</u>. For *F*_max_ and AUPRC, higher is better; for *S*_min_, lower is better.

| Method | MFO |  |  | BPO |  |  | CCO |  |  |
| --- | --- | --- | --- | --- | --- | --- | --- | --- | --- |
| | $F_{\max}$ | $S_{\min}$ | AUPRC | $F_{\max}$ | $S_{\min}$ | AUPRC | $F_{\max}$ | $S_{\min}$ | AUPRC |
| LLM | 0.603 | 7.175 | 0.609 | 0.440 | 36.593 | 0.417 | <u>0.706</u> | <u>6.813</u> | <u>0.662</u> |
| BPV | 0.666 | 6.434 | 0.575 | 0.465 | 41.715 | 0.410 | 0.692 | 7.068 | 0.554 |
| BPV + GNN | 0.667 | <u>6.082</u> | 0.638 | 0.463 | <u>35.932</u> | 0.438 | 0.691 | 7.004 | 0.589 |
| LLM + BPV | <u>0.688</u> | <u>6.082</u> | **0.662** | **0.486** | 41.715 | <u>0.446</u> | 0.700 | 7.004 | **0.742** |
| **ProtFun** | **0.693** | **5.930** | <u>0.658</u> | **0.486** | **35.478** | **0.455** | **0.710** | **6.796** | 0.597 |

## 4 Discussion

Identifying and elucidating the roles of proteins is crucial for comprehending biological processes. Detecting tens of thousands of protein functions in each of the BPO, MFO, and CCO categories is a complex multi-label classification task. In this study, we introduce a model that aims to improve protein function prediction by combining different data types. Specifically, we obtained protein sequence embeddings utilizing the pre-trained ANKH transformer LLM. We constructed a protein family network (PFN) using InterPro data and employed Graph Attention Networks (GAT) on this network to learn node embeddings. These embeddings were concatenated with the latent representations of protein signature data from InterPro to train a protein prediction model.

PPI networks provide a way to investigate the functional relationships resulting from the interactions between proteins. Employing GNN on PPI networks to predict protein functions is a commonly utilized strategy in literature and CAFA contests. Nevertheless, using PPI networks in this setting is subject to some constraints. First, PPI networks lack complete definition for each species, which could potentially restrict the ability of the model to generalize. Moreover, the interaction between two proteins does not indicate that they have identical functions. Although the presence of an interaction between two proteins suggests their participation in a shared biological process, it is possible for interacting proteins to have distinct biological roles (32). In this study, we present a more extensive network (i.e., protein family network) that considers supplementary biological characteristics of proteins, including their family structure, domain, and motif information. This methodology offers a more comprehensive and profound knowledge of protein functions and facilitates functional predictions that overcome the constraints of PPI networks.

The proposed approach offers several key advantages. First, it provides creation of a PFN by considering biological characteristics of proteins, including their family structure, domain, and motif information. Second, it processes embeddings obtained a protein LLM with GAT, which allows the model to effectively analyze both sequence- and networkbased data. Finally, by combining multiple data perspectives, such as sequence and binary protein representations acquired from InterPro, the hybrid model can leverage diverse data aspects, leading to a more nuanced understanding of protein features.

As a future work, to enhance the accuracy and speed of predictions in the model, we will explore using only the embeddings of functionally significant parts of proteins, such as domain regions, as input, rather than the complete protein sequences. By placing particular emphasis on functionally crucial protein sections can mitigate the risk of information loss caused by the length of the sequences and yield more precise predictions. Moreover, we will incorporate protein structure information into the model as features obtained from protein structures could improve protein function prediction.

## 5 Conclusion

In this study, we introduce a novel protein function prediction model, ProtFun. To facilitate integration of multi-modal datasets, we constructed a protein family network where node features were obtained from a protein LLM. Applying GAT on this network, we learned protein embeddings, which were concatenated with the latent representation of protein signatures from InterPro to train a protein function prediction model. Our results on the DeepGOZero, NetGO, and Deep-GraphGO datasets demonstrated that ProtFun outperformed the baseline and SOTA methods in predicting MFO, BPO, and CCO terms. We also performed ablation studies, which showed the importance of integrating diverse data modalities for effective protein function prediction.

## ACKNOWLEDGEMENTS

This work was supported by the National Institute of General Medical Sciences of the National Institutes of Health [R35GM133657], the National Artificial Intelligence Research Resource [NAIRR240068], and the startup funds from the University of North Texas.

