## Supplementary material for "ProtFun: A Protein Function Prediction Model Using Graph Attention Networks with a Protein Large Language Model": Table S1, Table S2,

### Supplementary Document

Table S1. Performance of the model on human, mouse, and *Arabidopsis thaliana* utilizing the DeepGOZero dataset.

| | $F_{max}$ (higher is better) | | | $S_{min}$ (lower is better) | | | **AUPRC** (higher is better) | | |
|---|---|---|---|---|---|---|---|---|---|
| Species (Taxon ID) | **MFO** | **BPO** | **CCO** | **MFO** | **BPO** | **CCO** | **MFO** | **BPO** | **CCO** |
| Human (9606) | 0.679 | 0.505 | 0.721 | 7.406 | 43.472 | 8.069 | 0.676 | 0.493 | 0.631 |
| Mouse (10090) | 0.673 | 0.449 | 0.691 | 6.491 | 52.155 | 7.203 | 0.667 | 0.425 | 0.531 |
| *Arabidopsis thaliana (3702)* | 0.782 | 0.461 | 0.750 | 3.377 | 24.795 | 5.434 | 0.777 | 0.420 | 0.631 |

Table S2. Performance comparison of the ProtFun model on CAFA species utilizing the NetGO dataset.

| | $F_{max}$ (higher is better) | | | $S_{min}$ (lower is better) | | | **AUPRC** (higher is better) | | |
|---|---|---|---|---|---|---|---|---|---|
| *Taxon ID* | **MFO** | **BPO** | **CCO** | **MFO** | **BPO** | **CCO** | **MFO** | **BPO** | **CCO** |
| 9606 | 0.799 | 0.531 | 0.696 | 2.722 | 23.185 | 6.676 | 0.805 | 0.495 | 0.617 |
| 3702 | 0.632 | 0.459 | 0.739 | 4.830 | 27.293 | 4.475 | 0.609 | 0.372 | 0.626 |
| 6239 | 0.735 | 0.442 | 0.723 | 4.814 | 33.092 | 3.184 | 0.769 | 0.396 | 0.607 |
| 7227 | 0.611 | 0.532 | 0.747 | 4.627 | 20.605 | 5.258 | 0.576 | 0.498 | 0.634 |
| 7955 | 0.671 | 0.363 | 0.706 | 5.831 | 25.936 | 6.107 | 0.626 | 0.290 | 0.433 |
| 9823 | 0.739 | 0.504 | 0.778 | 3.877 | 20.518 | 4.000 | 0.643 | 0.473 | 0.599 |
| 10090 | 0.627 | 0.410 | 0.659 | 5.381 | 44.610 | 10.717 | 0.615 | 0.363 | 0.556 |
| 10116 | 0.761 | 0.495 | 0.737 | 3.172 | 32.903 | 4.798 | 0.753 | 0.430 | 0.632 |
| 44689 | 0.747 | 0.392 | 0.705 | 3.462 | 16.158 | 5.516 | 0.652 | 0.316 | 0.637 |
| 284812 | 0.616 | 0.514 | 0.697 | 3.936 | 13.032 | 6.451 | 0.507 | 0.507 | 0.567 |
| 559292 | 0.640 | 0.441 | 0.751 | 3.350 | 21.991 | 6.325 | 0.460 | 0.425 | 0.639 |
